# GNAT: An Interactive Web Tool for Gene Neighbourhood Analysis

**DOI:** 10.1101/2025.11.03.686425

**Authors:** Nolen J. Timmerman, Malcolm Thakurdeen, Robert E.J. Wilson, Beatrice C.M. Fung, Brett Trost, Alan R. Davidson, Chantel N. Trost, Lingling Jin

## Abstract

Given the sequence of a protein, the Gene Neighbourhood Analysis Tool (GNAT) identifies homologues within microbial (bacterial, archaeal, or fungal) or viral databases, aligns and clusters their genomic neighbourhoods (GNs) by similarity, reports the taxonomic distribution of matched genomes, and generates interactive, publication-quality visualizations with customizable annotations, colours, and display options, along with categorical and quantitative summary analytics. GNAT’s intuitive interface enables a broad community of researchers to generate biological insights from GNs without requiring computational expertise and serves as a comprehensive and versatile platform for easily comparing GNs across the domains of life. GNAT is accessible at https://gnat.usask.ca.

## Introduction

Despite the exponential growth of sequence data in GenBank, which has doubled approximately every 18 months – outpacing Moore’s Law (1) – the functions of many genes remain unknown. One approach to infer gene function is to study its genomic context relative to other functionally characterized genes. This method is particularly powerful in prokaryotes and viruses, whose genomes contain operons, which are groups of genes controlled by a single promoter. These genes are transcribed together and then translated into discrete protein products that are often functionally related (2). Functionally related genes may also cluster in adjacent genomic locations outside of operons.

Well-characterized examples of gene clustering in prokaryotes include systems for pathogenicity/virulence (3), biosynthesis (4), and bacteriophage defense (5). In viruses, antidefense genes frequently cluster within phage genomes (6, 7). Gene clustering also occurs in eukaryotes, such as the penicillin biosynthetic cluster in fungi (8) and the human leukocyte antigen genes in humans (9). By studying genes within their immediate genomic contexts, we can gain insights into their functions, genomic organization, and evolutionary relationships.

Genomic neighbourhoods (GNs), which encompass the genes immediately upstream and downstream of a given gene, can be analyzed to identify clusters of functionally related genes. Moreover, by exploring GNs across multiple genomes, we can infer the potential functions of uncharacterized genes and detect patterns of gene conservation and organization across diverse species. GN analysis has been successfully used to discover more than 100 prokaryotic immune systems (5, 10, 11) and, in combination with additional comparative analyses, has revealed evolutionary links between prokaryotic and eukaryotic immune systems (12). Given the biological insights made possible by GN analyses, access to a simple, intuitive, and highly customizable bioinformatics tool that enables laboratory scientists to investigate GNs would be invaluable. Although numerous tools have been developed for this purpose (i.e. (13–19)), most have been designed to address the specific needs of individual research groups and, therefore, lack broad applicability. Many are further constrained by restrictive input formats, limited database options, inflexible search parameters, and rigid output configurations (Supplementary Table 1), which hamper customizability and overall utility. Some existing tools allow customization, such as the recent LoVis4u (19), but require familiarity with command-line interfaces and programming skills. Consequently, a truly accessible and user-friendly platform for the systematic exploration of GNs is still unavailable to most researchers.

To address this, we developed the Gene Neighbourhood Analysis Tool (GNAT), a user-friendly web application that enables researchers to search, analyze, visualize and compare GNs in an intuitive and highly customizable manner. GNAT requires only the amino acid sequence of a protein of interest as input, allowing researchers without computational or programming expertise to explore hundreds of GNs with ease. We anticipate that GNAT will expedite the discovery and characterization of novel genes and pathways, particularly in the fields of microbiology and immunology. Our major contributions include: 1) an interactive web application providing user-friendly access to the pipeline, several precomputed microbial (bacterial, archaeal, and fungal) and viral datasets, as well as comparative analysis features and customizable display options, 2) access to prokaryotic datasets with standardized, genome-based classification of bacterial and archaeal diversity, including integrated taxonomic and functional annotations for hypothesis-driven exploration, 3) two complementary protein annotation datasets: a ‘Bacterial Defense Systems’ dataset, consisting of anti-phage and other anti-mobile genetic element proteins; and, an ‘Anti-CRISPR Proteins’ dataset, consisting of CRISPR-Cas inhibitors and anti-CRISPR associated (Aca) proteins, and 4) the ability for research groups to build and maintain private, user-defined annotation libraries that can be reintegrated into the pipeline, enabling analyses tailored to their own datasets and research questions.

## Methodology

### Workflow Overview

GNAT is a bioinformatics pipeline written primarily in Bash and Python. Its core workflow consists of five main steps as outlined in Fig. 1a: 1) identifying distant orthologues via Position-Specific Iterative (PSI)-BLAST (20); 2) parsing the results and removing duplicates; 3) extracting genomic neighbourhoods; 4) clustering these neighbourhoods based on their similarity; 5) generating interactive visualizations for each cluster. A detailed description of our methodology is provided in the Supplementary Materials.

**Fig. 1.**
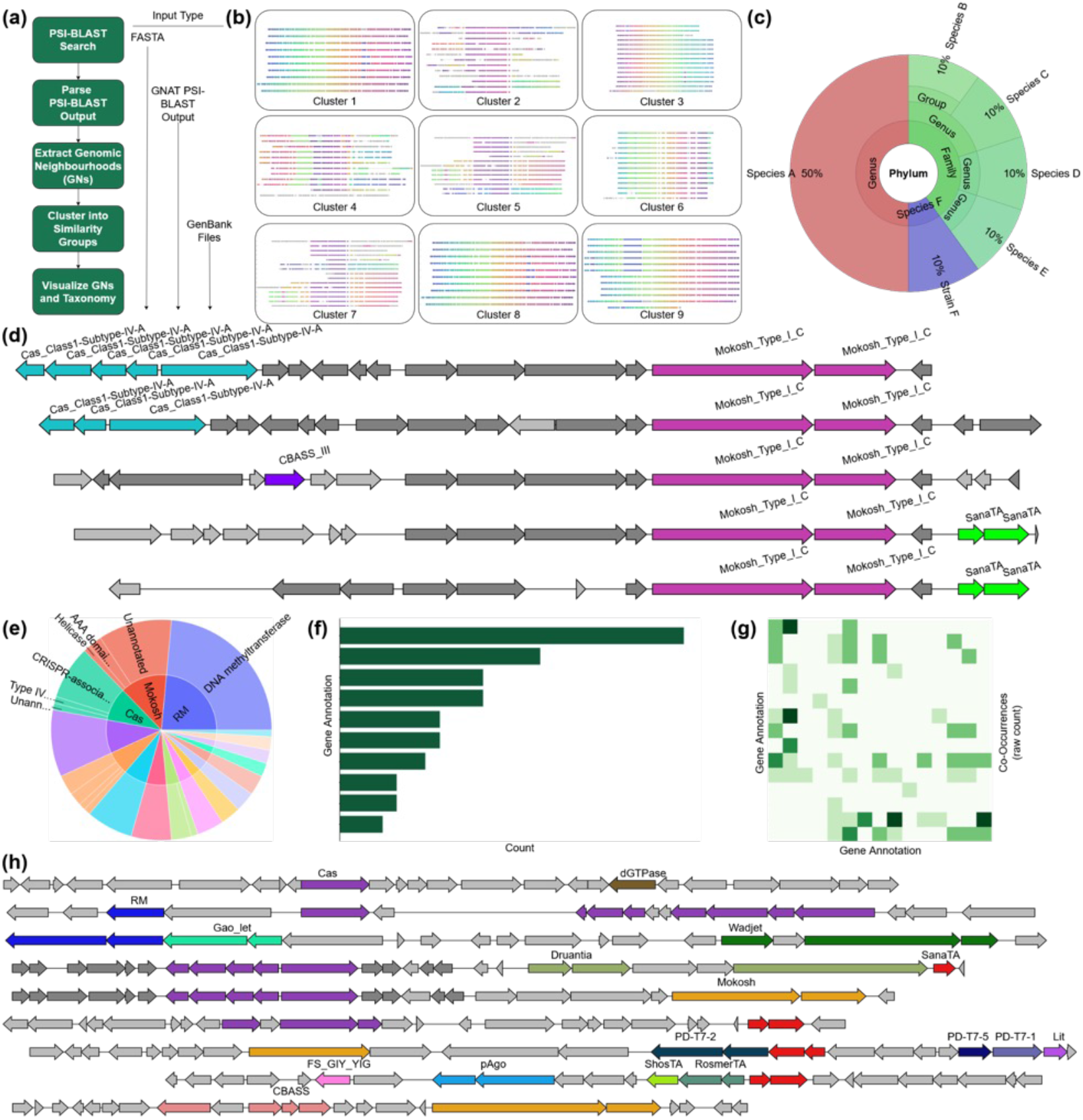
An overview of the GNAT workflow and its main outputs. (a) A simplified flowchart of the GNAT workflow. On the left are the main steps in the pipeline, and on the right are the input modes allowing users to enter the pipeline at various points. (b) The GNAT dashboard, presented to the user upon job completion, showing static images linking to each GN visualization. (c) A simplified taxonomic distribution plot summarizing the genomes matched. The default ‘collapsed’ mode is shown, which hides layers lacking deviations, allowing the ‘Family’, ‘Genus’, and ‘Species’ ranks to appear on the same layer. (d) A GN visualization generated with our modified version of clinker & clustermap.js. The query used to generate this plot was MokB (WP_227674162.1) of the Mokosh Type I-C bacterial defense system. The defense subsystem dataset and our alternate colouring mode were used to label and colour the plot. (e) A sunburst plot from the GNAT job summary showing the relative abundance of each defense system and its associated NCBI product annotations. (f) A bar chart from the GNAT job summary ranking the gene annotations detected in the generated GNs. (g) A heatmap from the GNAT job summary displaying the co-occurrence counts for the gene annotations. (h) A re-coloured GN visualization containing various co-localized defense systems, manually generated via the ‘GenBank Files’ input mode. The GNs in this visualization were pulled from the GNAT job summaries of SanaTA, Cas, and CBASS jobs.

### Input Modes

GNAT accepts three input modes with their relative entry points marked in Fig. 1a, providing a modular approach and the option to reuse previously generated results. The first input mode accepts a protein’s amino acid sequence in FASTA format as a PSI-BLAST query. The second uses existing PSI-BLAST results, allowing users to skip the homology search when re-clustering GNs. The third enables clustering of only selected GNs for targeted analysis, presentation, or publication.

### Homology Detection and GN Clustering (Steps 1-4)

In step 1, PSI-BLAST is used for homology searches (20). This iterative approach, though slower than standard protein BLAST (BLASTP) (21), provides enhanced sensitivity for detecting distant homologues. Step 2 parses the PSI-BLAST results and removes duplicate matches, which can arise when the same protein is identified in multiple iterations. Homologues are then filtered by percent identity (PI) and length to prioritize matches for GN extraction. A user-adjustable PI threshold (default = 30%) excludes distant hits that may not be biologically relevant to the user’s query. A user-adjustable length threshold (default = ± 20% relative to the query) is used to group proteins together (‘in-size’), while proteins outside this range are grouped separately (‘out-size’). This is a unique feature of GNAT, as current tools do not prevent GNs from proteins with partial or truncated regions or differing domain architectures from being clustered together, which can obscure biological interpretation. For instance, filtering by size allows one-domain and multi-domain proteins to form separate clusters, ensuring meaningful clustering.

In step 3, GNAT extracts the GNs surrounding each homologue. GNs consist of *n* genes (*n* = 15 by default) upstream and downstream of the homologue, resulting in GNs that contain up to 2*n* + 1 genes. In step 4, extracted GNs are filtered for redundancy and then clustered into groups of 5-15 genomes using an iterative *k*-means algorithm with 3-mer amino acid features (see Appendix for details). Clustering is performed for three size-filtered groups: in-size, out-size, and across all homologues. Dividing GNs into 5-15-genome clusters is required due to the computational limitations of the visualization software underlying GNAT, as well as to avoid lag and poor user experience when displaying more than 15 GNs on a standard user computer.

### Dashboard and Visualizations (Step 5)

Upon job completion, a dashboard is presented to the user (Fig. 1b), providing an overview of the run and enabling rapid identification of GN clusters of interest. By clicking on a static image in the dashboard, users are linked directly to the corresponding GN visualization. GNAT also outputs a plot summarizing the taxonomic diversity of the matched genomes, facilitating interpretation of evolutionary conservation and functional trends (Fig. 1c). For user convenience, all extracted GNs are output in GenBank format, allowing users to manually create visualizations that integrate up to 50 GNs from different queries using GNAT’s third input mode.

### Customization of GN Plots

#### Performance Optimizations

GNAT visualizations build on Clinker (13), extending its interactive features and streamlining the creation of publication-quality figures, while addressing key limitations—slow performance from computationally intensive global alignments for homologous GN detection and laggy interactivity caused by redundant gene-link data. GNAT’s clustering algorithm optimizes the underlying data structure, efficiently grouping homologous GNs without relying on global alignments and using carefully curated, non-redundant gene links. This enables smooth visualization of large datasets, allowing users to explore and interpret GN relationships without performance bottlenecks.

#### Interactive Visualization and Custom Labeling

We implemented several key customizations to automate annotation and enhance the exploration of GNs. These include customized labeling of all PSI-BLAST-matched genes, automatic orientation and alignment of GNs around them, and fine-tuned adjustments of scale, spacing, and other display parameters to optimize visualization of prokaryotic and viral genomes. Additionally, the legend displays the most prevalent NCBI product annotations for each homology group, helping users quickly infer each group’s biological function. Another interactive feature allows users to hover over legend groups to highlight corresponding orthologues in the plot, with the option to toggle among groups. A dynamic labeling option further allows the incorporation of label files from other analyses or merged outputs from prior runs, enabling users to update annotations without regenerating plots. Most importantly, the tier-labeling function supports custom label files containing additional annotation columns (e.g., defense system, defense subsystem, anti-defense system, anti-defense subsystem) and automatically recolours the plots accordingly. These customized annotations and colour schemes are illustrated in Fig. 1d. At every filtering step, we also generate interactive taxonomic distribution plots using the Krona tool (22), providing detailed insights into the evolutionary relationships among PSI-BLAST matches, similar to Fig. 1c.

To automate the annotation of GN visualizations, GNAT generates comma-delimited label files that can be easily viewed and edited in spreadsheet programs such as Microsoft Excel (see Appendix). Each label file initially contains a mapping between PSI-BLAST match protein accessions and the query protein annotation (taken from the first item in the FASTA header), along with the match e-value.

Existing label files can be uploaded to a new job, allowing GNAT to update match accessions if a new hit has a lower e-value, or to append accessions that are not already present. Users can further customize label files by adding additional columns for task-specific annotations. For instance, a ‘Defense System’ field can be added to the header, with accessions labeled accordingly (e.g., ‘Cas’) to denote defense-related genes (see Appendix). GNAT can also automatically generate such files when a user selects one of the provided label databases (see Appendix).

Plots annotated with these files can be recoloured using a drop-down menu to highlight groups according to the selected label tier, enabling customized visualizations. Figure 1d illustrates such a plot generated with a customized label file containing a ‘Defense Subsystem’ field.

### Summary Files and Analysis Outputs

After generating all GN visualizations, GNAT parses the annotation data to produce an HTML summary file containing key figures and statistics to facilitate GN analysis. Examples include: 1) a sunburst plot showing the relative abundance of each annotated protein and its associated NCBI product annotations (Fig. 1e), 2) a bar chart ranking the functional annotations detected in the generated GNs (Fig. 1f), and 3) a heatmap reflecting co-occurrence counts for gene annotations (Fig. 1g). Most figures in the GNAT summary file are derived from recorded co-occurrences of various annotations. The summary file also includes links to plots that contain an unusually high number of annotated genes. These outlier plots are identified using a linear regression model that predicts the expected number of annotated genes in each GN visualization based on the total number of genes and total number of GNs in the plot. Standardized residuals and approximate *p*-values are then computed, and plots with *p* ≤ 0.15 are highlighted as containing more annotations than expected.

### Web Application and Implementation

The GNAT web application is hosted on University of Saskatchewan servers using uWSGI and NGINX. The backend is implemented in Python with Flask, and the frontend is built using JavaScript, HTML, and CSS. GNAT is compatible with all major web browsers, with optimal performance on Google Chrome.

## Databases

### Overview

GNAT integrates three types of databases: genomic, taxonomy, and annotation. The genomic database, built from NCBI GenBank files, supports protein BLAST searches for homology detection and the extraction of corresponding GNs. The taxonomy database, an expanded version of that used by Krona, ensures all PSI-BLAST hits are mapped to their NCBI taxonomic IDs for representation in Krona plots. Finally, two annotation databases, which are essentially large label files, enable automatic GN annotation.

### Genomic

GenBank genome assembly summary files available on the NCBI FTP site (ftp://ftp.ncbi.nih.gov/genomes/genbank/) were used to construct most BLASTP databases. Genomes were downloaded as .gbff.gz files, from which all protein sequences were extracted to create a master FASTA file for BLASTP database construction. For larger databases exceeding PSI-BLAST query limits, the protein alias file (.pal) was divided into sets of 250 aliases, each queried separately. The corresponding GBFF files were retained for GN extraction based on PSI-BLAST results. Scripts enabling users to build custom databases from the NCBI FTP repository are available on the GNAT GitLab page.

GNAT provides BLASTP databases for bacterial, viral, and fungal genomes, including 20 genus-specific bacterial sub-databases representing the most abundant genera, as well as a combined bacterial–viral database. Additional GNAT-compatible BLASTP databases (GTDB bacterial, GTDB archaeal, and GTDB full) were also constructed from the Genome Taxonomy Database (GTDB) (23). A summary of all BLASTP databases is provided in Supplementary Table 2.

### Taxonomy

A taxonomy database is required to generate taxonomic distribution plots. While Krona creates a database mapping accessions to taxonomic IDs during installation, this database does not include protein sequences derived from whole-genome sequencing (WGS) projects. To cover the many sequences in GenBank originating from WGS data, we supplemented the Krona database with an additional protein mapping file, ensuring comprehensive taxonomic coverage.

### Annotation

GNAT allows users to select a ‘provided label database’ when submitting a job. If a label database is chosen, any genes in the user’s GNs that match entries in the selected database are annotated by appending extra columns to the user’s label file, providing additional functional or categorical information. The defense system dataset was obtained from Defense Finder (5) and expanded with accessions from the Identical Protein Group database from NCBI to improve coverage, as GNAT relies on protein accession numbers to match and annotate genes. Although not exhaustive, this combined dataset contains over 23 million proteins. An anti-CRISPR (Acr) protein database was built using entries from Bondy-Denomy *et al*. (24) as queries in PSI-BLAST searches against the combined bacterial–viral database, resulting in 144,619 annotated proteins. This broader coverage allows GNAT to detect and label hundreds of Acr and Aca proteins.

## Conclusion

Most genes in prokaryotic and viral genomes have unknown functions due to their extensive diversity. Yet, functionally related genes often occur together in operons or adjacent operons, enabling co-expression and horizontal exchange among bacteria and viruses. Thus, a practical way to infer protein functions, mechanisms of gene exchange, and evolutionary relationships is to analyze genomic neighbourhoods (GNs). Existing GN visualization tools often lack accessible interfaces, customizability, and analytical depth, limiting their use by non-bioinformaticians. GNAT addresses these limitations by enabling customizable searches against many databases, supporting built-in and user-defined annotations, and generating interactive GN visualizations and analytics. Starting with any protein of interest, homologues are identified and their GNs extracted. These neighbourhoods are aligned and clustered by similarity, and the taxonomic distribution of matched organisms is reported. The tool produces interactive, publication-quality visualizations with support for custom annotations, colours, and display options. With its user-friendly interface, GNAT enables a broad community of researchers to explore and interpret GN data.

